# Modulating DNA Polα Enhances Cell Reprogramming Across Species

**DOI:** 10.1101/2024.09.19.613993

**Authors:** Rajesh Ranjan, Binbin Ma, Ryan J. Gleason, Yijun Liao, Yingshan Bi, Brendon E. M. Davis, Guanghui Yang, Maggie Clark, Vikrant Mahajan, Madison Condon, Nichole A. Broderick, Xin Chen

## Abstract

As a fundamental biological process, DNA replication ensures the accurate copying of genetic information. However, the impact of this process on cellular plasticity in multicellular organisms remains elusive. Here, we find that reducing the level or activity of a replication component, DNA Polymerase α (Polα), facilitates cell reprogramming in diverse stem cell systems across species. In *Drosophila* male and female germline stem cell lineages, reducing Polα levels using heterozygotes significantly enhances fertility of both sexes, promoting reproductivity during aging without compromising their longevity. Consistently, in *C. elegans* the *pola* heterozygous hermaphrodites exhibit increased fertility without a reduction in lifespan, suggesting that this phenomenon is conserved. Moreover, in male germline and female intestinal stem cell lineages of *Drosophila*, *polα* heterozygotes exhibit increased resistance to tissue damage caused by genetic ablation or pathogen infection, leading to enhanced regeneration and improved survival during post-injury recovery, respectively. Additionally, fine tuning of an inhibitor to modulate Polα activity significantly enhances the efficiency of reprogramming human embryonic fibroblasts into induced pluripotent cells. Together, these findings unveil novel roles of a DNA replication component in regulating cellular reprogramming potential, and thus hold promise for promoting tissue health, facilitating post-injury rehabilitation, and enhancing healthspan.

## Introduction

The information encoded in DNA sequences constitutes the fundamental genetic material of life. DNA replication ensures the faithful copying of genetic information for reliable transmission to daughter cells during mitosis. In eukaryotic cells, DNA replication is tightly coupled with nucleosome assembly, which helps maintain or modify epigenetic information (*1–5*). Consequently, this process presents an opportunity for cells to prime for or initiate alterations in their identities (*6–8*). However, it remains unclear whether manipulating this process could affect cell fate determination during homeostasis or regeneration in multicellular organisms.

In living organisms, maintaining fitness requires the activity of adult stem cells to counteract cell loss during homeostasis and in response to injury. It has been hypothesized that somatic maintenance and reproduction have opposing effects on organismal lifespan (*9, 10*), although this relationship may be context-dependent, varying by sex, species, environment, and other factors (*11–14*). Tissue damage caused by pathogen infection or environmental changes rely on proper adult stem cell function to facilitate healing and growth. However, aging or injury often leads to the loss or impairment of stem cell activity (*15–24*), underscoring the need for effective strategies to enhance or reactivate adult stem cells for tissue maintenance or repair.

Additionally, induced pluripotent stem cells (iPSCs) (*25, 26*) offer an unprecedented avenue for regenerative medicine to treat various human diseases (*27, 28*). Researchers have been manipulating core transcription factors, signaling pathway components, microRNAs, and epigenetic modifications, among other approaches, to enhance iPSC reprogramming efficiency (*29–37*). The potential impact of fundamental biological processes, such as DNA replication, on reprogramming has started to be explored (*38–40*) but further studies are needed to fully understand their roles and mechanisms in multicellular organisms.

Here, we demonstrate that deliberately reducing Primase levels or DNA Polα activity enhances regenerative capabilities across various stem cell systems. In multiple stem cell models, progenitor cells in heterozygotes for genes encoding DNA Primase subunits exhibit enhanced reprogramming potential, leading to improved regenerative abilities during aging and tissue repair. These primed progenitor cells can functionally replace *bona fide* stem cells under both physiological and pathological conditions, suggesting a novel pathway for *in vivo* cell reprogramming within stem cell lineages. Furthermore, our *ex vivo* data using a well-established iPSC protocol show that an inhibitor modulating DNA Polymerase α (PolA1) facilitates the transition from differentiated cells to a pluripotent state more effectively.

## Results

### Male flies with reduced Polα levels have sustainable fertility during aging

*Drosophila* germline stem cell (GSC) lineages serve as a model system for studying stem cell maintenance, differentiation, and reprogramming (*41*). In both males and females, GSCs sustain the germline through asymmetric divisions, which produce a gonialblast (GB) in males (Fig. 1A, S1B) or a cystoblast (CB) in females (Fig. 2A). GBs and CBs subsequently undergo four additional mitotic divisions before transitioning into meiosis and differentiating into sperm or eggs.

**Figure 1:**
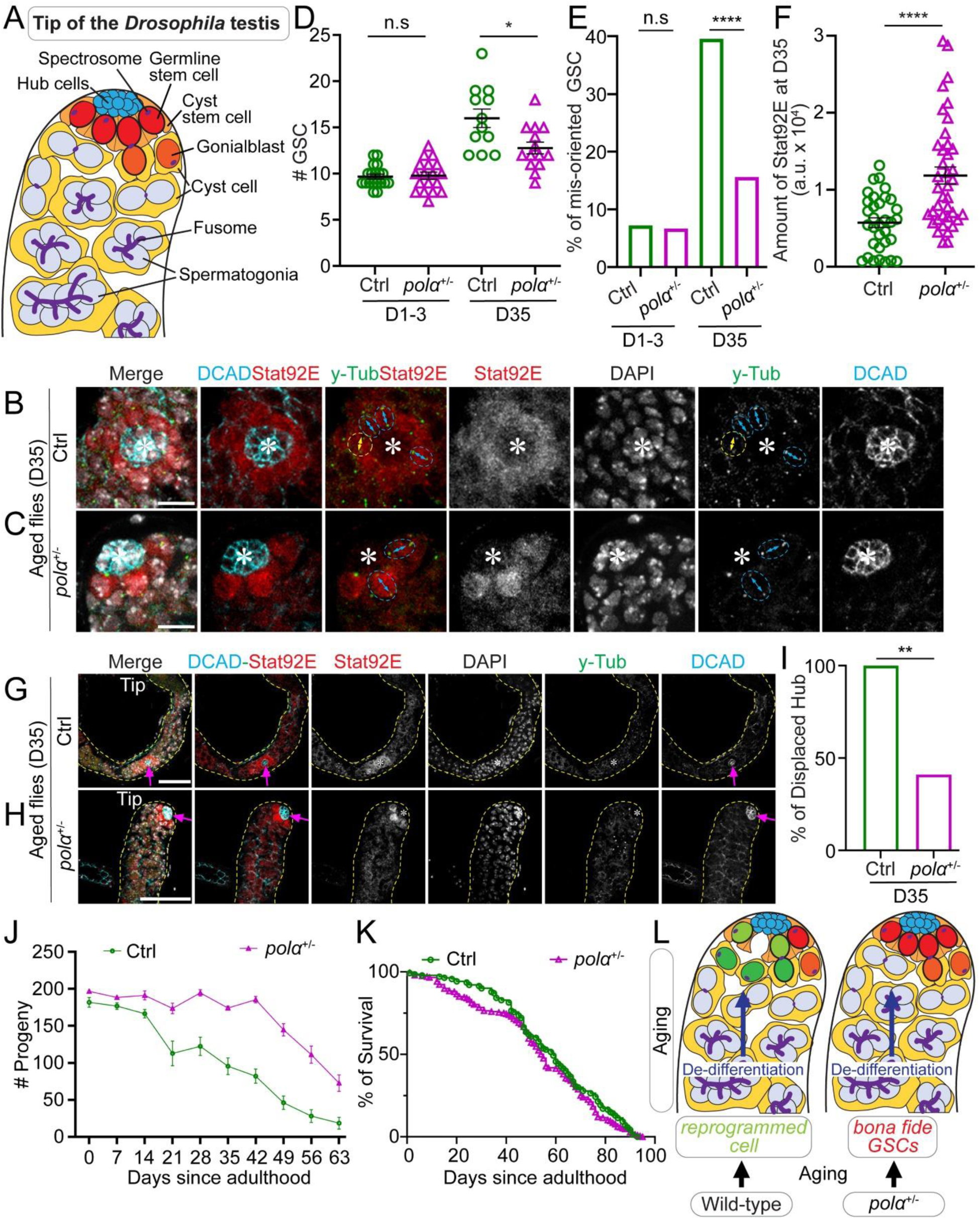
Male flies with compromised Polα have sustainable fertility during aging. (**A**) A cartoon depicting the apical tip of the *Drosophila* testis showing different cell types and their characteristic cellular features. (**B–C**) Representative images of the apical tips of 35-day old (D35) male testes, showing GSCs with oriented (cyan double-arrowed line pointing to the two separated centrosomes) and misoriented (yellow double-arrowed line) centrosomes in the control (Ctrl, **B**) and *polα50^+/-^*(*polα^+/-^*, **C**) flies. (**D**) Quantification of the number of GSCs in D1-3 and D35 Ctrl or *polα50^+/-^* testes [Ctrl D1-3 = 9.68 ± 0.27 (n = 19); *polα50^+/-^* D1-3 = 9.78 ± 0.39 (n = 18); Ctrl D35 = 16.00 ±1.0 (n = 12); *polα50^+/-^* D35 = 12.79 ±0.64 (n = 14)]. \**P*< 0.05, by Mann-Whitney test. (**E**) Quantification of the percentage of GSCs with misoriented centrosomes in D1-3 and D35 Ctrl or *polα50^+/-^* testes [Ctrl D1-3 = 7.19% (n = 153); *polα50^+/-^*D1-3 = 6.59% (n = 167); Ctrl D35 = 39.58% (n = 192); *polα50^+/-^*D35= 15.64% (n = 179)]. \*\*\*\**P* < 10^-4^, Chi-square test. (**F**) Quantification of the Stat92E immunostaining signals for both the Ctrl and *polα50^+/-^* GSCs in D35 testes. Ctrl GSC Stat92E= 5,746.94 ± 627.66 (n = 33) and *polα50^+/-^* GSC Stat92E= 11,872.31 ± 1,118.42 (n = 39), \*\*\*\**P* < 10^-4^ by Mann-Whitney test. (**G-H**) Representative images of the apical tips of D35 male testes, showing hub location, which is displaced from the testis tip in the Ctrl (**G**) but retained at the testis tip in the *polα50^+/-^* testes (**H**). Yellow dotted lines outline testes. (**I**) Quantification of the percentage of testes with displaced hub in D35 Ctrl testes (100%, n=14) or in the *polα50^+/-^*testes (40%, n=22). \*\**P* < 0.01, Chi-square test. (**J**) Quantification of fertility at different time points during aging (D0 to D63) for either the Ctrl (green line) or *polα50^+/-^* (magenta line) males (see Materials and Methods). Ctrl [D0 = 181.75 ± 6.65 (n = 16); D7 = 176.67 ± 4.46 (n = 18); D14 = 166.45 ± 5.74 (n = 20); D21 = 128.87 ± 15.67 (n = 18); D28 = 122.3 ± 12.38 (n = 20); D35 = 95.42 ± 11.01 (n = 22); D42 = 81.96 ± 9.37 (n = 29); D49 = 46.15 ± 9.15 (n = 13); D56 = 28.09 ± 8.71 (n = 11); D63 = 18.54 ± 8.02 (n = 13)]; and *polα50^+/-^* [D0 = 198.14 ± 4.90 (n = 22); D7 = 188.00 ± 4.83 (n = 21); D14 = 196.11 ± 17.35 (n = 22); D21 = 171.96 ± 8.02 (n = 26); D28 =198.05 ± 4.34 (n = 54); D35 =174.13 ± 3.21 (n = 22); D42 = 184.48 ± 6.16 (n = 45); D49 =139.87 ± 9.45 (n = 25); D56 = 46.15 ± 9.15 (n = 19); D63 = 46.15 ± 9.15 (n = 14)]. Two-way ANOVA (fixed-effects), for time factor: \*\*\*\**P*< 10^-4^; for genotype factor: \*\*\*\**P*< 10^-4^, for interaction: \*\*\*\**P*< 10^-4^. (**K**) The lifespan of control and *polα50^+/-^*flies. *P* > 0.99 by Long-rank (Mantel-Cox) test. (**L**) A model depicting the dedifferentiation and redifferentiation processes during aging. All values = Average ±SEM. Scale bar: 10μm (**B-C**), 50μm (**G-H**), hub: asterisk in (**B-C**) or arrow in (**G-H**).

**Figure 2:**
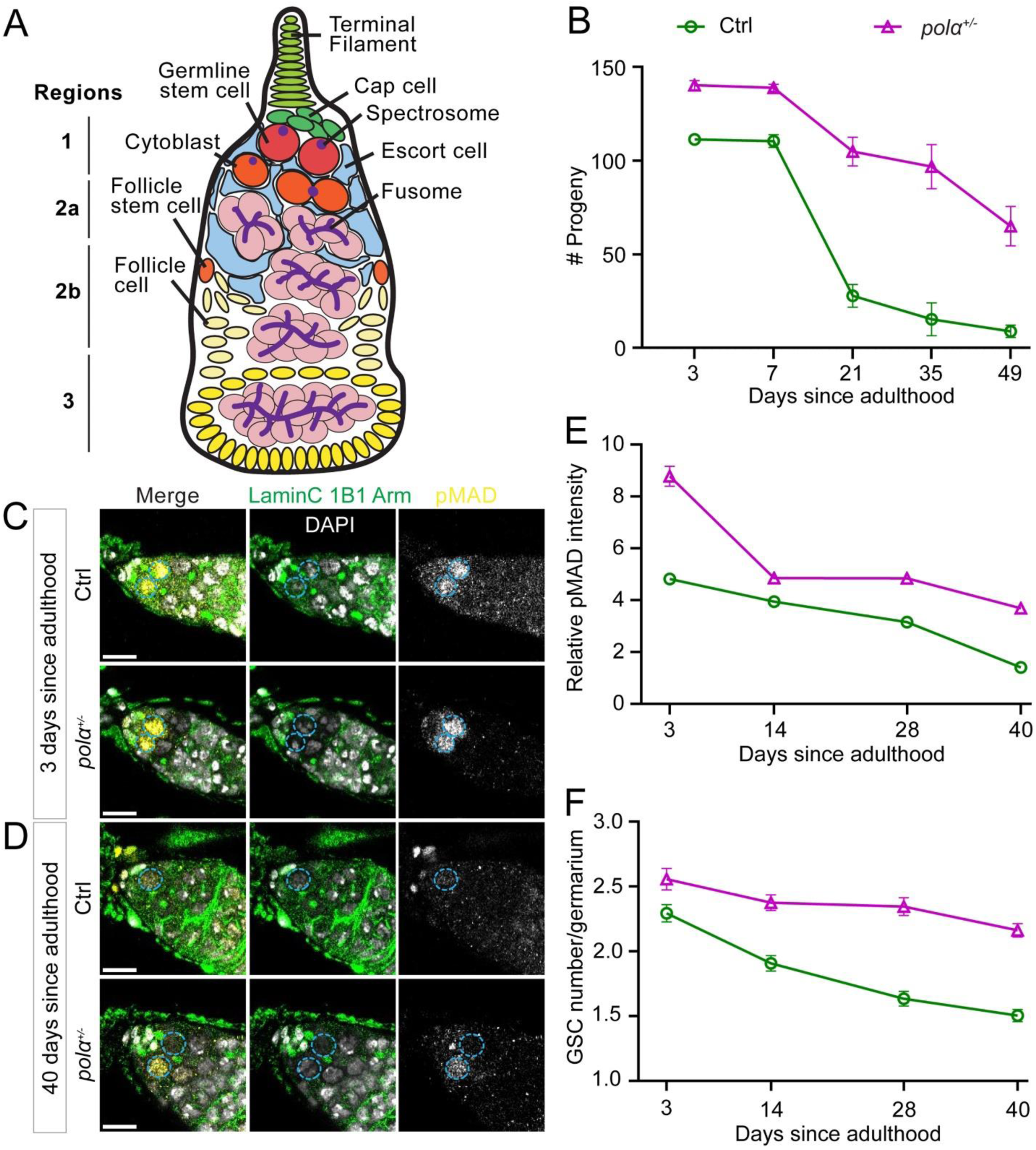
Female flies with reduced Polα levels have sustainable fertility during aging. (**A**) Illustration of *Drosophila* germarium, which can be divided into four regions: 1, 2a, 2b and 3. Each germarium contains 2-4 GSCs (red), locating at the most apical end of the germarium (Region 1). Each GSC divides into a new stem cell and a differentiating Cytoblast (orange). The Cytoblast undergoes 4 rounds of mitosis and form an egg chamber with follicle cells in Region 3, then leave the germarium for further oogenesis. (**B**) Quantification of female fertility at different time points during aging (D3 to D49) for either the Ctrl (green line) or *polα50^+/-^* (magenta line) females (see Materials and Methods). Two-way ANOVA (fixed-effects), for time factor: \*\*\*\**P*< 10^-4^; for genotype factor: \*\*\*\**P*< 10^-4^, for interaction: \*\*\*\**P*< 10^-4^. (**C-D**) Representative images of the germariums of 3-day old (**C**) and 40-day old (**D**) female, showing that GSC number is maintained in *polα*50^+/-^ females during aging, with higher pMAD signals than the control. GSCs are indicated by cyan dotted circle. Scale bar: 10μm. (**E**) Quantification of the pMAD immunostaining signals for both the Ctrl and *polα50^+/-^*GSCs during aging (see Materials and Methods). Ctrl [D3= 4.822 ± 0.1764 (n = 201); D14= 3.949 ± 0.1847 (n = 185); D28= 3.155 ± 0.1949 (n = 167); D40= 1.405 ± 0.08970 (n = 193)] and *polα50^+/-^* [D3= 8.785 ± 0.3895 (n = 118); D14= 4.850 ± 0.1747 (n = 251); D28= 4.845 ± 0.1653 (n = 237); D40= 3.684 ± 0.1423 (n = 377)]. Two-way ANOVA (fixed-effects), for time factor: \*\*\*\**P*< 10^-4^; for genotype factor: \*\*\*\**P*< 10^-4^, for interaction: \*\*\*\**P*< 10^-4^. (**F**) Quantification GSC number for both the Ctrl and *polα50^+/-^* females during aging. Ctrl [D3= 2.294 ± 0.06658 (n = 85); D14= 1.907 ± 0.06057 (n = 97); D28= 1.634 ± 0.05755 (n = 101); D40= 1.504 ± 0.04736 (n = 127)] and *polα50^+/-^* [D3=2.558 ± 0.08354 (n = 43); D14= 2.375 ± 06139 (n = 104); D28= 2.346 ± 0.06800 (n = 104); D40= 2.163 ± 0.05197 (n = 172)]. Two-way ANOVA (fixed-effects), for time factor: \*\*\*\**P*< 10^-4^; for genotype factor: \*\*\*\**P*< 10^-4^, for interaction: \*\**P*< 0.01. All values = Average ±SEM.

Previous studies have shown that aging or injury often leads to the loss of GSCs or their activity (*15–24, 42*). In the male germline, progenitor spermatogonial cells (SGs) can "dedifferentiate" to re-enter the stem cell niche (*17–19, 21, 43*). This process could occur both under physiological conditions like aging (*18, 44*) or following genetic ablation of GSCs (*17, 19, 43, 45*). Typically, the centrosomes of GSCs are oriented perpendicularly to the GSC-niche interface to facilitate mitotic spindle formation (*46–48*). In contrast, dedifferentiated GSC-like cells often carry misoriented centrosomes. This misorientation could activate a "centrosome orientation checkpoint," which blocks successful mitosis and arrests GSCs in the G2 phase until corrected (*18, 49, 50*). As GSCs gradually turnover with age, the niche is increasingly occupied by dedifferentiated SGs, potentially leading to dysfunctional germline and reduced fertility in aged males (*18*).

In a previous study, GSCs exhibit reduced levels of several DNA replication machinery components compared to SGs, including DNA Polymerase α (Polα) (*51*). This suggests that DNA replication machinery may critically regulate germline function in a stage-specific manner, potentially influencing key processes such as differentiation and dedifferentiation. In our study, we investigated whether reducing Polα in SGs could enhance their ability to restore GSC activities upon dedifferentiation. Since DNA replication is essential for cell cycle progression and survival, we aimed to modulate replication machinery components without inducing lethality or cell cycle arrest. To this end, we used a null allele of the *polα50* gene, which encodes DNA Primase Subunit 1 (Prim1) involved primarily in lagging strand synthesis. The heterozygous *pola50^+/-^*flies are viable and exhibit no readily apparent phenotypes. We then evaluated whether *pola50^+/-^* SGs can more effectively replace GSCs compared to controls by examining cellular features and redifferentiation capabilities of the dedifferentiated GSC-like cells, with a focus on the effects of aging.

In this experimental design, both control and *polα50^+/-^* male flies were examined in parallel from newly eclosed (D0) to 63 days old (D63, Fig. S1A). The control flies were obtained by crossing two wild-type strains (Materials and Methods) to avoid any potential advantages from outcrossing, which was also used to generate the *polα50^+/-^*heterozygotes. We first assessed GSC cellular features and testicular morphology in males from different age groups. Consistent with previous reports (*18, 52, 53*), the number of GSCs in D1-3 control and *polα50^+/-^* males was comparable (Fig. 1D). By D35, the GSC number showed a slight difference, with control testes having an average of three more GSCs than *polα50^+/-^*males (Fig. 1B-D). However, the percentages of GSCs with misoriented centrosomes changed more dramatically: at D1-3, the two genotypes showed no difference, but by D35, this percentage in control males was approximately 2.5 times higher than in *polα50^+/-^* males (Fig. 1B-C, 1E). Additionally, the critical "stemness" transcription factor Stat92E (*54–57*) was significantly elevated in *polα50^+/-^* GSCs compared to control GSCs in D35 testes (Fig. 1B-C, 1F). Morphologically, testes from D35 control males often had abnormal niche anatomy, such as a displaced hub structure (Fig. 1A, 1G, 1I, S1C-E) and a thin overall testicular structure with elongating and mature sperm in the terminal differentiating regions (Fig. 1G, S1F). In contrast, testes from D35 *polα50^+/-^* males displayed a normal hub structure at the testis apical tip (Fig. 1A, 1H-I) and a normal testicular structure with properly organized terminal differentiating regions (Fig. 1H, S1G). The displaced hub in older control flies may result from age-dependent decreases in adhesion molecule expression (*16*). This anatomical shift of the hub structure from the apical tip to the center of the testis may contribute to the increased number of GSCs, which were counted as the germ cells surrounding the hub (Fig. 1A, 1D, 1G, 1I).

To further assess the redifferentiation capabilities of dedifferentiated GSCs during aging, we conducted a time-course fertility assay using males from different age groups. Intriguingly, *polα50^+/-^* males maintained persistent fertility from D0 to D63, with a noticeable reduction only beginning at D42 (Fig. 1J). Even after D49, *polα50^+/-^* males still retained relatively high fertility until D63, whereas the control males were almost sterile by this time. In stark contrast, control males exhibited a continuous decline in fertility from D14 to D63 (Fig. 1J). These results demonstrate a pronounced divergence in male fertility between *polα50^+/-^* and control males across different stages of aging (Fig. 1L, S1H).

### Female flies with reduced Polα levels have sustainable fertility during aging

*Drosophila* oogenesis relies on the activity of female GSCs [Fig. 2A, (*58*); reviewed in (*59, 60*)]. In the female GSC niche, Bone Morphogenetic Protein (BMP) signaling from the niche regulates GSC identity and activity by phosphorylating Mothers against decapentaplegic (pMad), which in turn activates Daughters against decapentaplegic (Dad) expression for GSC maintenance and function (*61–70*). We next investigated whether the enhanced *pola50^+/-^* male germline activity can also be detected in the female germline. As previously shown, aging induces multiple changes in the female germline, including decreased GSC number, reduced BMP signaling (*65, 71*), and deteriorated female fertility over time (*72*).

We first tested the fertility of *pola50^+/-^* female flies over a 49-day period. We found that *pola50^+/-^* females consistently displayed higher fertility compared to control females from D3 to D49 (Fig. 2B). Although *pola50^+/-^* females showed reduced fertility at D49, their fertility was still significantly higher than that of D49 control females and even exceeded that of D21 control females. In contrast, control females were almost completely sterile by D35 (Fig. 2B). These results demonstrate that *pola50^+/-^* flies have sustained fertility in both males and females.

To explore the underlying mechanisms contributing to the enhanced female fertility, we first investigated BMP signaling activity in female GSCs. Immunostaining experiments revealed consistently higher levels of pMad in *pola50^+/-^* GSCs compared to control GSCs over a 40-day period (Fig. 2C-E). Sustained pMad levels led to consistently higher GSC numbers in *pola50^+/-^* ovaries than in control ovaries, with significant differences in GSC numbers evident throughout the 40-day period (Fig. 2F). Taken together, these findings indicate that GSCs in *pola50^+/-^* female flies exhibit substantially higher GSC maintenance, elevated levels of "stemness" factors, and increased germline activity during aging.

Remarkably, in a longevity assay [adapted from (*73*), Materials and Methods], both *pola50^+/-^* and control flies exhibited comparable lifespans (Fig. 1K). Notably, there were no significant differences in lifespan between control and *pola50^+/-^* flies, regardless of whether males, females (individual sex data not shown), or both sexes combined were considered (Fig. 1K). These findings confirm that the sustained fertility observed in *pola50^+/-^* males and females does not come at the expense of a shortened lifespan, indicating genuine reproductive longevity and extended fertility lifespan.

### *C. elegans* with reduced Polα levels have higher fertility

In *C. elegans*, the *pola-1* gene encodes the homolog of the POLA1 catalytic subunit of the DNA Polα–Primase complex. During adulthood, germline POLA-1 plays a critical role in GSC maintenance and germ cell proliferation (*74*). Here, we investigated whether *pola-1* heterozygosity could enhance hermaphrodite fertility. We used a null allele of the *pola-1* gene, *pola-1(gk5576)* (*75*). The *pola-1^+/-^* heterozygous strain is viable and shows no discernible phenotypic abnormalities.

We first compared the fertility of *pola-1^+/-^* heterozygotes to wild-type hermaphrodites. Interestingly, a significantly larger brood size in *pola-1^+/-^* worms was observed compared to wild-type controls (Fig. 3B). To further explore whether *pola-1^+/-^* heterozygotes exhibit higher GSC maintenance during aging, we used a GFP reporter (*76*) that specifically labels the GSC and progenitor cells in the gonad (Fig. 3A). Live-imaging of this reporter in *pola-1^+/-^* heterozygotes and wild-type worms revealed that reducing POLA-1 levels confer advantages to a higher GSC number compared to wild-type later into adulthood (D5 in Fig. 3C-D). These results suggest that, as observed in *Drosophila*, reduction of POLA-1 levels *via* heterozygosity significantly enhances fertility and GSC maintenance in *C. elegans*.

**Figure 3:**
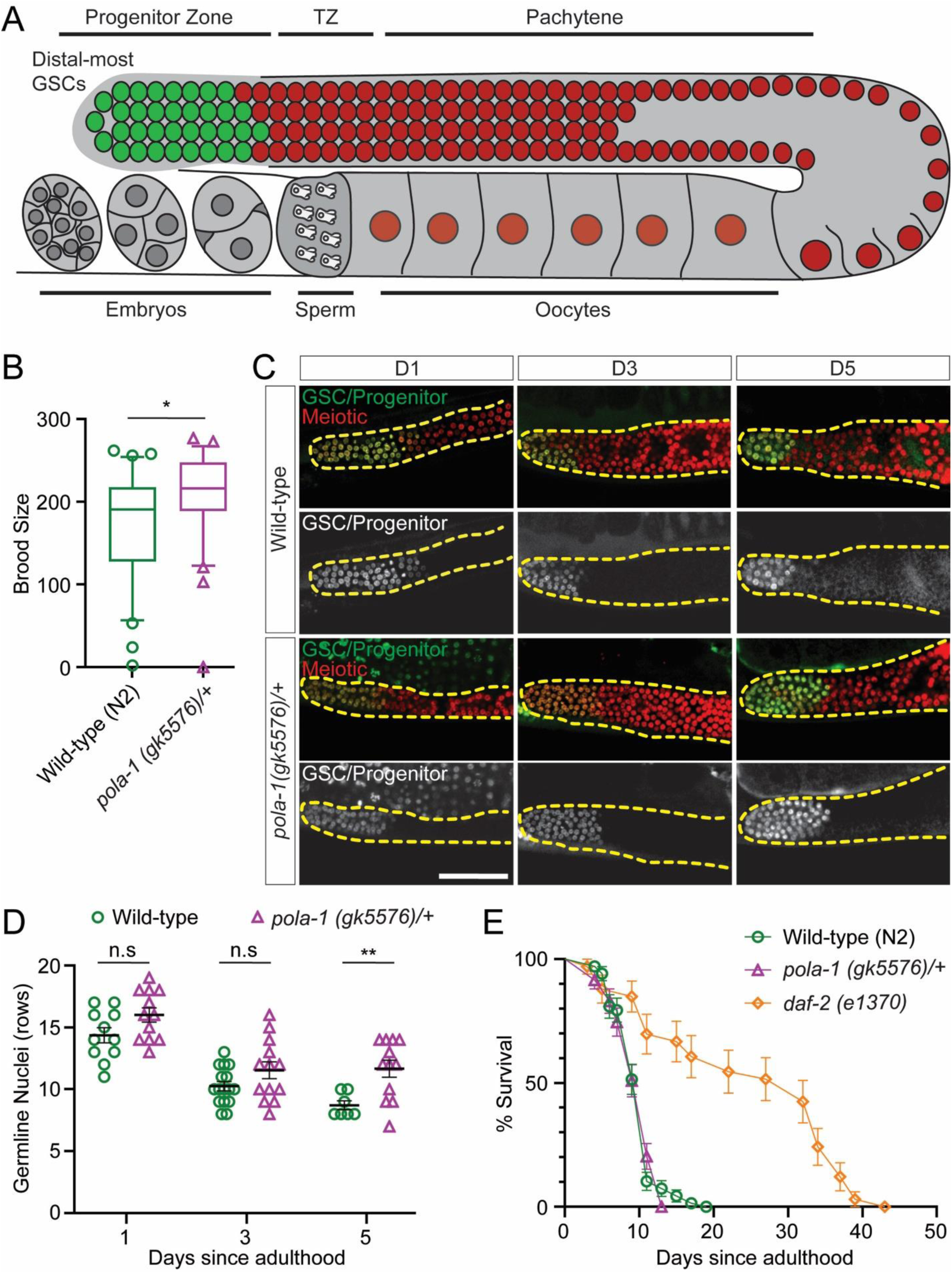
*C. elegans* with reduced POLA1 levels have higher fertility. (**A**) Illustration of the *C. elegans* hermaphrodite gonad (*110, 111*). Germline nuclei are arranged in a spatiotemporal pattern progressing from the distal most GSCs [green: GFP under the control of the *pie-1* promoter and the *zif-1* 3’UTR (*76*); red: mCherry labeling all germline nuclei]. The transition zone (TZ) contains meiotic leptotene/zygotene stages. Oocytes become fully cellularized by late diakinesis, pass through the spermatheca filled with sperm and undergo early embryonic development in utero. (**B**) Brood sizes for *pola-1(gk5576)/+* (n=30) and *wild-type* (n=30). Each data point represents the number of living larvae from individual worms with the corresponding genotype. \**P*< 0.05 by unpaired t test. (**C**) Representative images of *wild-type* and *pola-1(gk5576)/+* taken at Day 1, Day 3, or Day 5 of adulthood. The strain GC1413 rrf-1[pk1417; naSi2 (Pmex5::H2B::mCherry::nos-2 3’UTR); teIs113 (Ppie-1::GFP::H2B::zif-1 3’UTR)] was used. The dashed lines outline the gonads. (**D**) Quantification measured by rows of cells from the distal end. All values = Average ±SEM. *P* = 0.06 [Day 1, *pola-1(gk5576)/+* (n= 12), *wild-type* (n=11)]; *P* = 0.09 [Day 3, *pola-1(gk5576)/+* (n= 13), *wild-type* (n=16)]; \*\**P*< 0.01 [Day 5, *pola-1(gk5576)/+* (n= 12), *wild-type* (n=7)]; by unpaired t test. (**E**) Survival analysis of *wild-type*, *pola-1(gk5576)/+*, and *daf-2(e1370)*. No significant difference between *wild-type* and *pola-1(gk5576)/+*, by Long-rank (Mantel-Cox) test. Scale bar, 50 μm.

Next, we assessed whether the observed increase in fertility of *pola-1^+/-^*heterozygotes affects their lifespan. Here, we included three strains: wild-type (N2), *pola-1^+/-^* heterozygotes, and the well-characterized long-lived *daf-2(e1370)* mutant (*77*). We found no significant difference in the lifespan between *pola-1^+/-^* heterozygous and the wild-type strains (Fig. 3E). In contrast, the *daf-2* mutants nearly doubled their lifespan relative to wild-type worms, consistent with the previous report (*77*). Together, these results indicate that reduced POLA-1 levels enhance *C. elegans* fertility without compromising lifespan, suggesting that this phenomenon is conserved between flies and worms.

### Male flies with reduced Polα levels have enhanced regeneration capabilities in testes

In addition to declining stem cell activity during aging, stem cells often demonstrate regenerative capabilities in response to tissue damage. For example, genetic ablation of *bona fide* GSCs can trigger dedifferentiation of the progenitor SGs in the *Drosophila* male germline (*43, 45, 53*). This acute approach can complement studies of gradual changes that accumulate over time during aging.

To achieve this goal, we studied how GSCs recover after genetic ablation. We ectopically expressed the *grim* gene (*78, 79*) in early-stage germ cells using a temporally controllable system (*nos-Gal4ΔVP16*; *tubulin-Gal80^ts^*; *UAS-grim*), followed by a recovery period to allow dedifferentiation (Fig. 4A-B). Grim inhibits apoptotic antagonists and promotes apoptosis (*80–82*).

**Figure 4:**
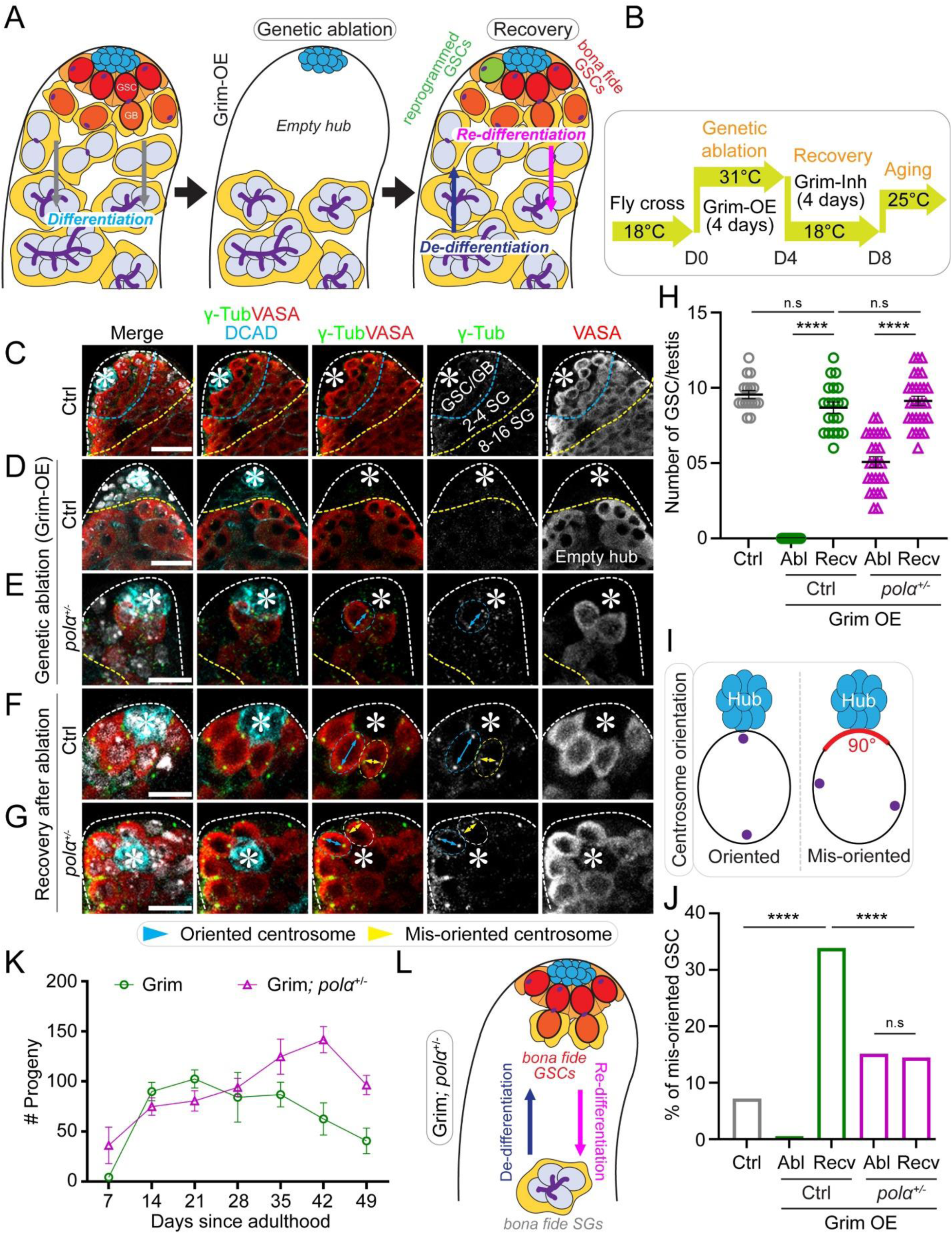
Dedifferentiated spermatogonial cells with reduced Polα levels tend to have properly oriented centrosomes and re-differentiate for increased male fertility. (**A**) A cartoon depicting the genetic ablation by overexpression of Grim (Grim-OE) that results in depletion of early-stage germ cells followed by recovery. (**B**) The temperature shift scheme to activate Grim expression (at 31°C, Grim-OE) followed by inhibiting Grim expression (at 18°C, Grim-Inh) using the Gal4; Gal80^ts^ system in adult flies. (**C**) A control (Ctrl) testis tip showing early-staged germ cells. (**D**) A representative image of the control (Ctrl, *UAS-grim*) testis tip showing a complete ablation of early-staged germ cells upon Grim-OE for four days at 31°C. (**E**) A representative image of the *polα50^+/-^*; *UAS-grim* testis tip showing significantly reduced GSCs number upon Grim-OE for four days at 31°C. (**F-G**) Upon recovery for four days at 18°C, a representative image of the Ctrl *UAS-grim* (**F**) and a representative image of the *polα50^+/-^*; *UAS-grim* (**G**) testes tips showing GSCs with oriented centrosomes (cyan double-arrowed line pointing to the two centrosomes) and GSCs with misoriented centrosomes (yellow double-arrowed line pointing to the two centrosomes), respectively. In (**C-G**): White dotted outlines indicate the testis tip, cyan dotted lines demarcate the GSC/GB region, from cyan to yellow dotted lines denote the 2-4 cell SGs, beyond yellow dotted line is the 8-16 cell SG region. (**H**) Quantification of the number of GSCs per testis in the Ctrl [Ctrl = 9.56 ± 0.27 (n=16)] and Grim-OE testes after genetic ablation [Abl (Ctrl)= 0 ± 0 (n=22)] and upon recovery [Recv (Ctrl)= 8.7 ± 0.36 (n=20)]; in the *polα50^+/-^*; *UAS-grim* after genetic ablation [Abl (*polα^+/-^*)= 5.08 ± 0.36 (n=26)] and upon recovery [Recv (*polα^+/-^*)= 9.14 ± 0.32 (n=28)]; \*\*\*\**P* < 10^-4^, Mann Whitney test. (**I**) A cartoon depicting the quantification of oriented *versus* misoriented centrosomes in GSCs. (**J**) Quantification of the percentage of GSCs with misoriented centrosome in the Ctrl without genetic ablation (Ctrl= 7.19%, n = 153), in the Ctrl *UAS-grim* after genetic ablation [Abl (Ctrl)= 0%, n= 22] and upon recovery [Recv (Ctrl)= 33.91%, n=174]; in the *polα50^+/-^*; *UAS-grim* after genetic ablation [Abl (*polα^+/-^*)= 15.16%, n=132] and upon recovery [Recv (*polα^+/-^*)= 14.45%, n=256]. \*\*\*\**P* < 10^-4^, Chi-square test. (**K**) Quantification of fertility at different time points during genetic ablation (D0 to D4 at 31°C) followed by recovery (D4 to D8 at 18°C) and aging (D8 to D49 at 25°C) for either the Ctrl (green line) or *polα50^+/-^*(magenta line) males with Grim overexpression (see **B** and Materials and Methods). Grim [D7 = 4.34 ± 2.61 (n = 41); D14 = 89.88 ± 8.80 (n = 43); D21 = 102.43 ± 8.58 (n = 51); D28 = 84.09 ± 24.90 (n = 11); D35 = 86.81 ± 12.35 (n = 32); D42 = 62.5 ± 16.68 (n = 14); D49 = 40.64 ± 12.85 (n = 11)]; and Grim; *polα50^+/-^* [D7 = 36.08 ± 18.19 (n = 12); D14 = 74.92 ± 8.71 (n = 25); D21 = 80.55 ± 10.21 (n = 20); D28 = 94.11 ± 9.11 (n = 35); D35 = 124.68 ± 17.24 (n = 22); D42 = 141.68 ± 13.02 (n = 25); D49 = 96.42 ± 9.59 (n = 24)]. Two-way ANOVA (fixed-effects), for time factor: \*\*\**P*< 10^-3^; for genotype factor: \*\*\*\**P*< 10^-4^, for interaction: \*\*\**P*< 10^-3^. (**L**) A cartoon depicting recovery from genetic ablation, a model to investigate de-differentiation and re-differentiation. Scale bar: 25μm (**C-D**), 10μm (**E-G**), asterisk: hub.

This genetic ablation approach effectively caused GSC death in both control (Fig. 4C-D) and *polα50^+/-^*testes (Fig. 4E), as evidenced by the substantially decreased GSCs following *grim* overexpression (Fig. 4H). The increased retention of GSCs in *polα50^+/-^* testes compared to controls may result from higher resistance to Grim overexpression or faster dedifferentiation. Nevertheless, during recovery, dedifferentiation led to significantly increased GSC-like cells in both control (Fig. 4F, 4H) and *polα50^+/-^* (Fig. 4G-H) testes. However, dedifferentiated GSCs often exhibited misoriented centrosomes (Fig. 4F-G, 4I), consistent with previous reports (*18, 52, 53*).

Intriguingly, we found that dedifferentiated GSCs in *polα50^+/-^* testes had significantly fewer instances of misoriented centrosomes compared to control testes (Fig. 4J). Notably, despite the nearly doubling of GSC numbers after recovery in *polα50^+/-^* testes (Fig. 4H), the percentage of GSCs with misoriented centrosomes remained unchanged (Fig. 4J), indicating that nearly all dedifferentiated *polα50^+/-^*GSCs tended to have properly oriented centrosomes. These results demonstrate that dedifferentiated GSCs in *polα50^+/-^*testes possess more *bona fide* GSC-like cellular features, such as properly oriented centrosomes, in addition to molecular features, such as Stat92E expression (Fig. 1F), compared to dedifferentiated GSCs from control testes.

To further assess the redifferentiation capabilities of dedifferentiated GSCs following genetic ablation, fertility tests were conducted using both control and *polα50^+/-^* males after recovery from *grim* overexpression (Fig. 4B). During the initial phase after recovery (D7 to D28), both genotypes exhibited comparable fertility (Fig. 4K). However, starting from D35, *polα50^+/-^* males showed significantly higher fertility than the controls, with this difference persisting up to D49 (Fig. 4K). These data demonstrate that in the male germline of *polα50^+/-^* flies, dedifferentiated GSCs can more effectively replace *bona fide* GSCs compared to controls (Fig. 4L), suggesting enhanced regenerative potential.

In summary, our results demonstrate that dedifferentiated *pola50^+/-^* GSCs can effectively replace *bona fide* GSCs under both physiological (Fig. 1) and pathological (Fig. 4) conditions. This suggests that fine-tuning the activity of specific DNA replication components, such as Primase, may be sufficient to restore GSC-like features in dedifferentiated cells during aging and tissue repair. Given the conserved role of DNA replication in cellular reprogramming, targeting specific DNA polymerases could represent a promising strategy in regenerative medicine.

### Female flies with reduced Polα levels have enhanced regeneration capabilities in midgut

In addition to the germline maintained by GSCs, *Drosophila* intestinal stem cells (ISCs) in the midgut represent a model somatic stem cell system (Fig. 5A-B), characterized by a well-defined lineage, abundant and easily distinguishable ISCs, as well as responsiveness to environmental cues such as nutrition, infection, and aging (*83–87*). We asked whether the enhanced regeneration abilities observed in the male germline are preserved in this somatic stem cell lineage.

**Figure 5:**
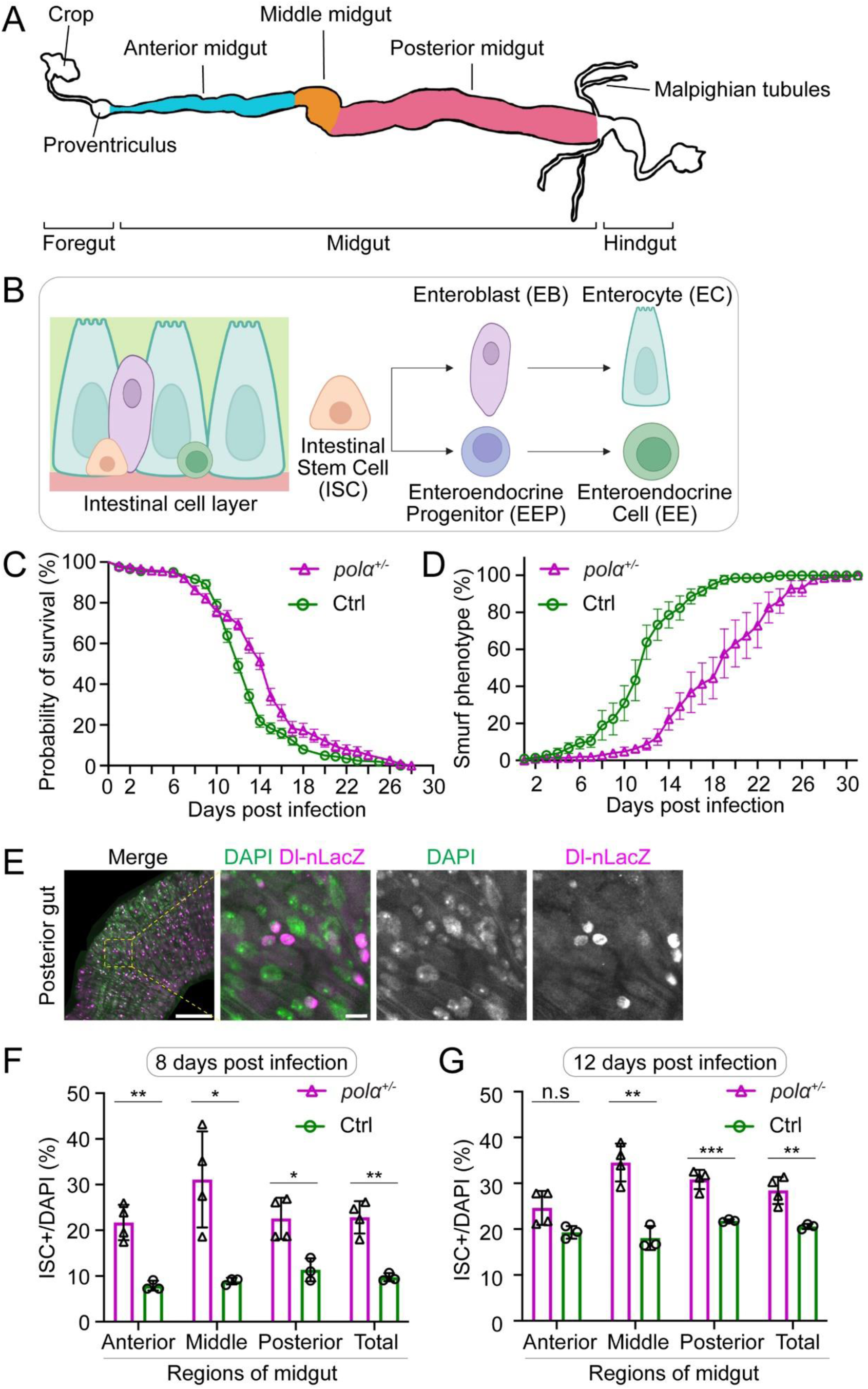
The *polα50* heterozygotes have enhanced regeneration capabilities in midgut. (**A**) *Drosophila* midgut anatomy: The midgut, located between the Proventriculus (Cardia) and Malpighian tubules, is divided into anterior, middle, and posterior sections based on morphology. (**B**) The fly midgut has a defined stem cell lineage performing diverse functions. Intestinal Stem Cells (ISCs), located at the basal membrane, are multipotent and self-renewing. ISCs differentiate into Enteroblasts (EBs) or Enteroendocrine Progenitor Cells (EEPs). EBs give rise to Enterocytes (ECs), responsible for secreting digestive enzymes and absorbing nutrients, while EEPs develop into Enteroendocrine Cells (EEs), which regulate hormonal balance through gut hormone secretion. (**C**) Probability of survival post-infection: Control (Ctrl, green line, n= 202) *vs. pola50^+/-^* heterozygotes (magenta line, n= 268). The median survival time: Ctrl= D12 *vs. pola50^+/-^*= D15. Kaplan-Meier test: Log-rank (Mantel-Cox) test, *P*< 10^-4^. (**D**) Smurf assay post-infection: Control (Ctrl, green line, n= 210) *vs. pola50^+/-^*heterozygotes (magenta line, n= 210). Two-way ANOVA (mixed-effects), for time factor: \*\*\*\**P*< 10^-4^; for genotype factor: *P*< 10^-3^. (**E**) Immunostaining of midguts: Delta-nlacZ (magenta), DAPI (green). Scale bar: 100μm for the zoom-out and 10μm for the zoom-in images. (**F-G**) Percentage of ISCs across different midgut regions of Ctrl (n= 3) and *pola50^+/-^* heterozygotes (n= 4) on Day 8 (**F**) and Day 12 (**G**) post-infection: The percentage of ISCs are significantly higher (Unpaired t test) across different regions compared to the control on both time points, ns: not significant (*P*> 0.05), * *P*< 0.05, \*\**P*< 10^-2^, \*\*\**P*< 10^-3^.

To determine whether *pola50^+/-^* heterozygotes confer advantages in response to tissue damage, we infected flies with a lethal strain of *Chromobacterium subtsugae*, a bacterium that is orally toxic to various insects, including *Drosophila melanogaster*, and used as an insecticide (*88, 89*). The *C. subtsugae* strain *ΔvioS* causes lethality of flies through producing secondary metabolites, which are normally inhibited by the VioS repressor and regulated by the quorum sensing system (*89*).

To validate the lethal effect of the *ΔvioS* strain, we implemented a regime of a brief starvation followed by feeding flies with either Luria-Bertani (LB) media (non-infected group) or *ΔvioS*-containing media (infected group). Both groups were then monitored for survival on fresh fly food over time (Fig. S2A). The non-infected control flies exhibited consistently high survival rates throughout a 28-day time course (Fig. S2C). In stark contrast, the infected control flies showed a sharp decline in survival starting from Day 8 (D8), with near-zero survival by day 20 post-infection (Fig. 5C). In a parallel assay using *pola50^+/-^* heterozygotes, the non-infected group showed almost no difference from control flies (Fig. S2C), while the infected group exhibited improved survival compared to controls, marked by a delayed decline in survival and a significantly extended post-infection lifespan in *pola50^+/-^* heterozygous flies (Fig. 5C).

To investigate the mechanism of oral pathogen-induced lethality, we introduced a blue dye into the food (referred to as ’smurf food’) and fed it to the flies post-infection (*90*) (Fig. S2A). This assay allows visualization of intestinal damage. In healthy flies, the dye remains confined to the intestine (Fig. S2B). However, pathogen infection causes increased intestinal permeability, resulting in blue pigment leakage into the body cavity, which can be visualized and quantified as a “smurf phenotype” (Fig. S2B). Both non-infected control and *pola50^+/-^* flies showed very low percentages of flies with the smurf phenotype (Fig. S2D). Contrastingly, infected *pola50^+/-^* flies consistently showed a lower percentage of smurf phenotype than infected control flies during a 30-day period (Fig. 5D). These results demonstrate improved maintenance in the midgut of *pola50^+/-^* heterozygotes compared to controls in response to pathogen-induced tissue damage.

To further understand tissue repair at the cellular level, we examined the expression pattern of Delta, a key ISC stem cell fate determinant (*91, 92*). Using a *Delta*-nuclear lacZ (*Dl*-nLacZ) reporter to mark ISCs (*93–95*), we quantified the percentages of the nLacZ-positive cells at three distinct regions of the midgut (Fig. 5A) at two time points (D8 and D12 post-infection) in both *pola50^+/-^* and control flies. These timepoints were chosen to correspond with the onset of rapid decline (D8) and approximately 50% survival (D12) of post-infected control flies (Fig. 5C). We found that ISCs were consistently higher in *pola50^+/-^* than in control flies across different midgut regions, resulting in significantly increased ISCs in *pola50^+/-^* midguts compared to the control at both timepoints (Fig. 5E-G), suggesting higher self-renewal or maintenance abilities of *pola50^+/-^*ISCs compared to the control. In summary, *pola50^+/-^* flies show increased resistance to pathogen-induced tissue damage in the midgut, leading to improved survival during post-injury recovery.

### Slightly reducing Polα activity enhances the efficacy of human iPSC induction

Thus far, we have demonstrated that compromising one allele of the Polα- or Primase-encoding genes in flies and worms can enhance the sustainability of adult stem cell systems or increase their resilience to tissue damage. Next, we aimed to explore whether pharmacologically modulating Polα activity could yield similar benefits, potentially establishing it as a viable therapeutic strategy for promoting stem cell regeneration.

Using human neonatal dermal fibroblasts, we conducted induced pluripotent stem cell (iPSC) reprogramming experiments with the CytoTune™-iPS 2.0 Sendai Reprogramming Kit (*96*), which overexpresses the transcription factors OCT3/4, SOX2, KLF4, and C-MYC (*25*). We employed a Polα inhibitor that impedes the DNA binding affinity and primer elongation activity of the DNA Polymerase α subunit 1 (PolA1) (*97*). Given that this inhibitor blocks DNA replication at relatively high concentrations (0.1 -1 µM in Fig. S3A-B), we first optimized its concentration to minimize any adverse effects on DNA replication itself, as indicated by comparable S-phase indices between treated and untreated fibroblasts (Fig. S3A-B). Subsequently, we used an optimized concentration of the PolA1 inhibitor during the iPSC induction process (10-20nM).

Using the PolA1 inhibitor at the optimized concentration (Fig. 6A), we observed that reducing Polα activity during the early reprogramming stage enhances this process, as indicated by the earlier appearance, higher rate, and larger size of iPSC colonies in inhibitor-treated cells compared to untreated cells at the same timepoints during reprogramming. Cells treated with the PolA1 inhibitor showed higher ratios of SSEA-4-positive cells, an early pluripotency marker (*98, 99*), starting from D6 and extending to D13 (Fig. 6B-C). Intriguingly, the well-established ’stemness’ marker NANOG was detectable as early as D2 in PolA1 inhibitor-treated cells and remained more abundant compared to the control from D2 to D13 (Fig. 6D-E, S3C). Meanwhile, the fibroblast differentiation marker Platelet-Derived Growth Factor Receptor alpha (PDGFRα) was less enriched in the PolA1 inhibitor-treated cells compared to the control from D2 to D13 (Fig. 6D-E, S3C). Furthermore, PolA1 inhibitor-treated cells exhibited brighter staining for TRA-1-60, a live cell dye for ’stemness’ (*100*), compared to the control cells at D11 (Fig. S3D).

**Figure 6:**
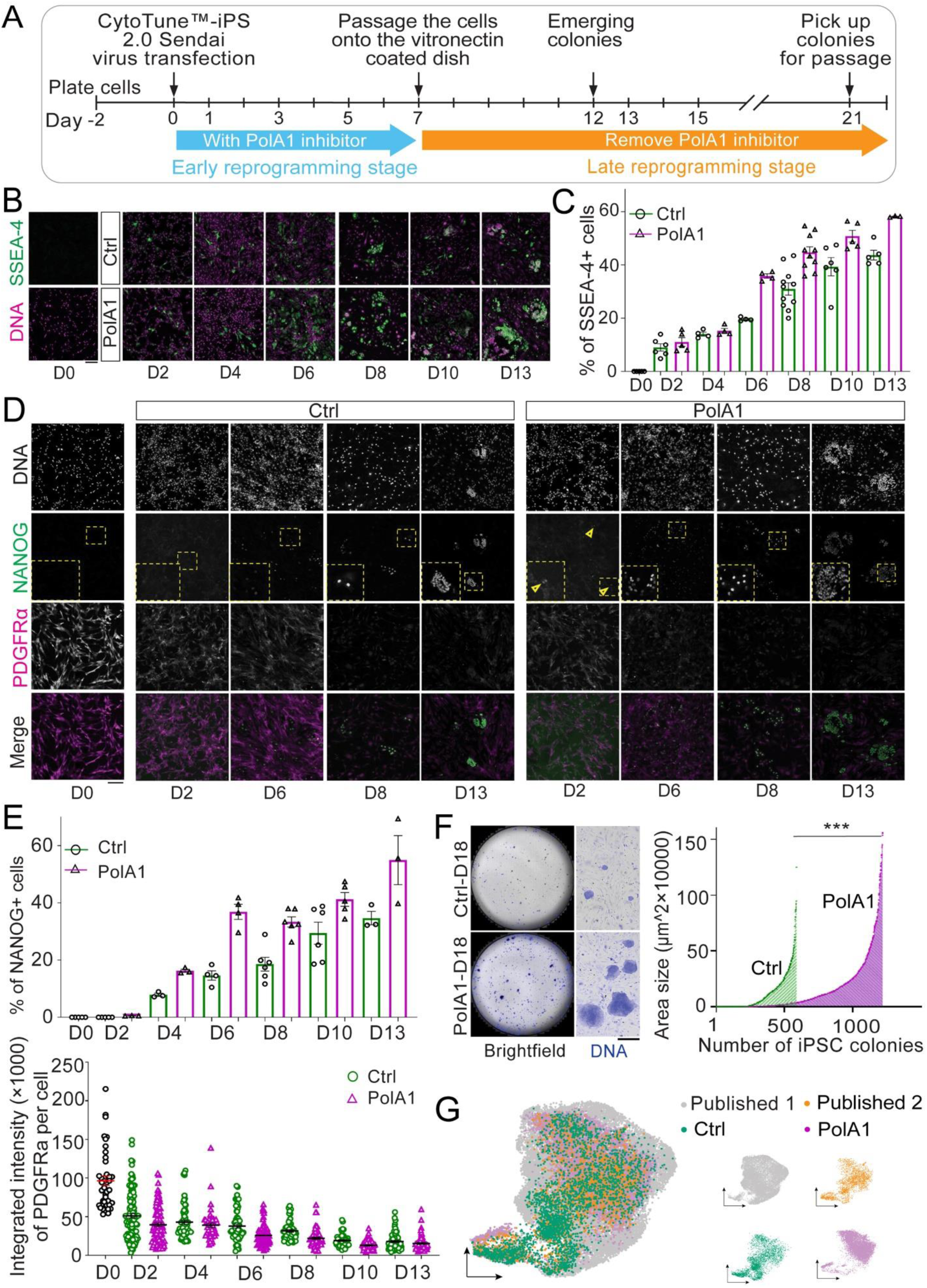
PolA1 inhibition enhances the human iPSCs reprogramming efficiency. (**A**) The workflow of CytoTune™-iPS 2.0 Sendai virus reprogramming used to generate human iPSCs with and without PolA1 inhibitor. (**B**) The SSEA-4 staining and (**C**) The percentage of SSEA-4+ cells at different time points during human iPSC reprogramming. Two-way ANOVA (fixed-effects), for time factor: \*\*\*\**P*< 10^-4^; for inhibitor factor: **** *P*< 10^-4^; for interaction: \*\**P*< 0.01. (**D**) The NANOG and PDGFRα staining and (**E**) The percentage of NANOG+ cells and integrated intensity of PDGFRα staining signals per cell at different time points during human iPSC reprogramming. NANOG+ cells: two-way ANOVA (fixed-effects), for time factor: \*\*\*\**P*< 10^-4^; for inhibitor factor: \*\*\*\**P*< 10^-4^; for interaction: \**P*< 0.05. PDGFRα intensity: two-way ANOVA (fixed-effects), for time factor: \*\*\*\**P*< 10^-4^; for inhibitor factor: \*\*\*\**P*< 10^-4^; for interaction: *P*= 0.158 (not significant, ns). (**F**) The whole dish imaging of human iPSCs stained with DNA dye Hoechst 33342 with and without PolA1 inhibitor. More and larger iPSC colonies were detected in PolA1 inhibitor-treated cells at D18 after reprogramming induction. \*\*\**P* <10^-3^, by Mann Whitney test. (**G**) snRNA-seq data of human iPSCs with or without PolA1 inhibitor during induction are visualized in UMAP, together with two publicly available scRNA-seq datasets of human iPSCs. Seurat was used to generate the plot after removing the batch effect with its “CCAIntegration” method.

Finally, DNA staining revealed a higher incidence and larger colony size in PolA1 inhibitor-treated cells compared to control cells at D18 (Fig. 6F). Together, these results indicate a better transition from differentiated to pluripotent state in the PolA1 inhibitor-treated cells.

To further investigate the properties of the reprogrammed iPSCs, immunostaining for well-known stem cell markers, including the transcription factors NANOG, OCT4, and SOX2, validated the pluripotent state of human iPSCs derived from both PolA1 inhibitor-treated and control cells at the end of the reprogramming processes (Fig. S3E). To assess the quality of the resulting iPSCs, we performed single-nucleus RNA sequencing (snRNA-seq) and compared reprogrammed iPSCs generated in the presence and absence of the PolA1 inhibitor with published data (*101, 102*). Our analysis showed that the obtained iPSCs exhibited strong pluripotency markers and a transcriptome profile largely overlapping with those of well-established human iPSCs (Fig. 6G, S3F). These findings confirmed that the iPSCs we generated maintain molecular features comparable to well-characterized iPSCs and that treatment with the PolA1 inhibitor did not compromise iPSC quality. In conclusion, our results suggest that a selective and slight inhibition of PolA1 can enhance the generation of human iPSCs, improving both the efficiency and dynamics of reprogramming.

## Discussion

The loss of stem cells or their functionality is often exacerbated by aging or injury, prompting the need for regenerative strategies *in vivo*. In this study, we found that manipulating a key DNA replication component allows non-stem cells to effectively substitute *bona fide* stem cells, thereby maintaining stem cell activity. Importantly, these dedifferentiated cells can re-differentiate into functional progeny cells, ensuring sustained lineage functionality. A low dose of the Polα inhibitor, targeting the lagging strand PolA1, enhances cellular reprogramming, further underscoring its potential in promoting regenerative processes. Therefore, our findings hold promise for application in other adult stem cells and in inducing pluripotent cells from patient-derived cells.

Reducing Primase levels or Polα activity, either through heterozygosity or low concentrations of an inhibitor, could shift the epigenetic symmetry typically present in progenitor cells (*51*), potentially initiating epigenetic asymmetry as a ’priming’ step to potentiate cell fate changes. However, these calibrated conditions may not be fully deterministic for altering cell fates. Certain molecular features, such as the transcriptome, may remain unchanged or show only minor alterations, allowing progenitor cells to undergo normal differentiation under non-challenging conditions, such as homeostasis in young adults. Conversely, under conditions that require natural or induced cell reprogramming, such as aging or tissue damage, these potentiated cells might adopt a partially reprogrammed state, enabling them to outperform control cells in both dedifferentiation and re-differentiation processes *in vivo*.

Recent studies have shown that manipulating epigenetic asymmetry can lead to defects in mouse embryonic stem cell differentiation (*103, 104*) and mouse development (*105*). These experiments use mutations in replication components with histone chaperoning activities, such as the *mcm2-2A* or *pole3* mutants, which normally function as DNA helicase and leading-strand DNA polymerase, respectively. Our experimental design differs in several key aspects from these conditions. First, *mcm2-2A* or *pole3* mutants induce a high degree of histone incorporation asymmetry across the genome (*106–108*), which can be detrimental to normal cell functions, including stem cell differentiation. In contrast, we used *primase* or *polα* heterozygotes to reduce the levels of the full-length protein or applied low concentrations of the PolA1 inhibitor to slightly reduce its activity. These conditions may be sufficient to shift the balance from epigenomic symmetry to asymmetry without causing global changes. Alternatively, they may change the speed of replication forks. Notably, it has been demonstrated that DNA replication speed can influence cell fate decisions in early mouse embryogenesis, with a slower replication process promoting cellular reprogramming toward totipotency (*109*). Second, the molecular mechanisms underlying embryonic stem cell differentiation *in vitro* could differ from those involved in dedifferentiation *in vivo* or iPSC reprogramming *in vitro*. Nevertheless, our findings underscore the need for further studies to better understand the regulation of these processes, which are essential for normal development and hold promise for regenerative medicine.

In summary, our results suggest an intriguing new approach to stimulate stem cell regeneration in response to injury or aging-induced stem cell depletion, thereby promoting reproductive longevity, healthy aging, and an increased healthspan.

## Data availability

GEO accession number for the snRNA-seq data is GSE277293 (released on 09/15/2027 or earlier).

## Acknowledgments

We thank Drs. John Kim, Keji Zhao, Caitlin Pozmanter, Jingchao Zhang and Chen lab members for helpful suggestions. We thank Drs. Yukiko Yamashita, Erika Matunis, Allan Spradling and Jane Hubbard for fly and worm strains used in this study. We thank Johns Hopkins Integrated Imaging Center for confocal imaging. Supported by NIH 5T32GM007231 (Y.B., B.D.), NIH R35 GM127075 and R01 HD102474, and the Howard Hughes Medical Institute (X.C.).

## Author Contributions

R.R., B.M., R.G., Y.L., Y.B., B.D., G.Y., N.B. and X.C. conceptualized the study. R.R., B.M., R.G., Y.L., Y.B., B.D., G.Y., M.C., V.M., and M.C. performed all experiments and data analyses. R.R., B.M., R.G., Y.L., Y.B., B.D., G.Y., N.B. and X.C. wrote the manuscript.

## Supplemental Information

### Materials and Methods

#### *Drosophila* strains and husbandry

Fly strains were raised on standard Bloomington media. All flies were raised at 25°C unless noted otherwise. The following fly strains were used: *polα50* P-element insertion (BL-27205) (*51*), *nos-Gal4 (with VP16)* on the 2^nd^ chromosome (*112*), *nos-Gal4* (*without VP16* or *DVP16*) on the 2^nd^ chromosome [(from Yukiko Yamashita, Whitehead Institute, USA) and used in (*113*)], *Delta-nuclear lacZ* reporter on the 3^rd^ chromosome (Dr. Allan Spradling, Carnegie Institute, USA). The *UAS-grim* flies (*78*) (from Erika Matunis, Johns Hopkins School of Medicine, USA) were crossed with *nanos-Gal4DVP16; tubulin-Gal80^ts^* for genetic ablation experiments to induce dedifferentiation (see below).

All experiments using the *pola50^+/-^* were outcrossing the *pola50*/Balancer stock to a wild-type stock (*y,w*) to have the *pola50* P-element insertion allele over a wild-type chromosome. To avoid any potential effects brought by outcrossing, control flies were from outcrossing two wild-type strains: Oregon-R and *y,w*.

#### *C. elegans* strain maintenance

*C. elegans* were maintained on nematode growth medium agar plates using Escherichia coli OP50 as a food source and cultured according to standard methods (*114*). The following strains were used in this study: N2, VC4505 [*pola-1(gk5576)III/+* heterozygotes], CB1370 [*daf-2(e1370) III*], GC1413 rrf-1[*pk1417; naSi2 (Pmex5::H2B::mCherry::nos-2 3’UTR); teIs113 (Ppie-1::GFP::H2B::zif-1 3’UTR)*] (from Jane Hubbard, New York University, USA).

#### Immunostaining

Immunostaining experiments were performed using standard procedure for *Drosophila* testes (*115*), ovaries (*116*) and midguts (*117*). All secondary antibodies were the Alexa Fluor-conjugated series (1:1,000; Molecular Probes).

For immunostaining of *Drosophila* testes, primary antibodies used were DE-cadherin (10:200; DHSB AB_528120), γ-Tubulin (1:200; Sigma-Aldrich AB_T6557), Stat92E (1:500; from Denise Montell, University of Santa Barbara, CA, USA), VASA (1:500; from Ruth Lehmann, Whitehead Institute, USA), Armadillo (1:100; DSHB N2 7A1), Traffic Jam (1:100, from Mark Van Doren, Johns Hopkins University, USA), anti-H4K20me2/3 (1:400; Abcam ab78517), and anti-H3S10ph (1:2000; Cell Signaling Technology 9701).

For immunostaining of *Drosophila* ovaries, ovaries were fixed in 4% formaldehyde in 0.3% PBST (1x PBS, 0.3% Triton X-100), washed twice for 10 min in 0.3% PBST, and blocked in 5% normal goat serum (NGS, Jackson ImmunoResearch lab, 005-000-121) in 1% PBST (1x PBS, 0.1% Triton X-100) overnight, followed by a two-day primary antibody incubation at 4°C (diluted in 0.3% PBST with 5% NGS). Then, ovaries were washed three times for 20 min in 0.3% PBST and twice for 30 min in 0.3% PSBT with 5% NGS, and incubated with secondary antibodies for 2 hours (diluted in 0.3% PBST with 5% NGS) at room temperature. In the last 30 min of secondary incubation, Hoechst (Thermo Fisher Scientific, 33342) was added. Last, they were washed three times for 20 min in 0.3% PBST before mounting in Vectashield (Vector Laboratories H100010). Primary antibodies include pMAD (1:800; Cell Signaling Technology 9516), Hts (1:20; DSHB, 1B1), α-spectrin (1:50; DSHB, 3A9), Armadillo (1:50; DSHB, N2 7A1), and LaminC (1:100; DSHB, LC28.26).

For Immunostaining of *Drosophila* midgut, fly intestines were dissected in pre-chilled Schneider’s media within 30 minutes, followed by fixation with 4% formaldehyde in PBST (1X PBS+0.1% TritonX-100) for 1 hour at room temperature on a nutator. Wash tissues in PBST for 10 minutes, repeating twice and perform an additional wash for 30 minutes. Block in 5% NGS + 1% BSA (Bovine Serum lbumin) for at least two hours at room temperature or overnight at 4°. Primary antibodies include β-Galactosidase (1:500; Abcam ab9361), H3T3P (1:500; Millipore Sigma 05-746R), diluted in PBST +5% NGS + 1% BSA for overnight incubation at 4°. Wash tissues in PBST for 10 minutes, repeating twice and perform an additional wash for 30 minutes. After secondary antibody incubation with secondary antibodies (diluted in PBST +5% NGS + 1% BSA) for 2 hours at RT. Wash tissues in PBST for 10 minutes, repeating twice and perform an additional wash in PBST for 30 minutes. Remove the last wash and mount the tissue with Fluoromount-G™ Mounting Medium with DAPI (Invitrogen,00-4959-52). The stitched images were then converted into Imaris file format (Imaris 10.1 (RRID:SCR_007370) and the number of both the nLacZ positive cells and DAPI were counted using the built-in spot analysis on Imaris. We then manually checked the cell count to filter out any false positive and false negative signals mis-detected by the software.

#### Quantification of pMAD immunostaining signals

To measure pMAD signals, z-stacks (sum of slices) were generated at 0.5 μm intervals of individual germarium. Using the draw tool in Fiji, a circle indicating the GSC nuclei based on Hoechst staining was drawn and the mean fluorescence intensity (MFI) was taken. Another circle was drawn in Region 2b and the mean fluorescence intensity was taken as background (BG). Stemness Index was defined as (MFI - BG)/BG to eliminate batch difference. GSCs were identified and quantified, based on the spectrosome morphology and position: round α-spectrin signals at the anterior tip of the germarium and next to cap cells.

#### Tile scan imaging of the *Drosophila* gut

To examine regeneration ability during gut infection, we conducted tile scan imaging of the entire gut with high spatial resolution. For this, adult *Drosophila* intestines were immunestained and mounted on slides for imaging. All tile scans were performed using a spinning disk confocal microscope equipped with two qCMOS QUEST cameras (Hamamatsu), an X-Light V3 confocal spinning disk system, a high precision Piezo XY stage, and a LDI-laser module. Images were acquired using a 63x Olympus oil object. The VisiView software (BioVision) was used to outline the gut, set the Z-stack, select individual lasers, and acquire images with 2×2 binning. After acquiring individual tile scans, VisiView software was also used to stitch them together, reconstructing the entire gut. The images were processed using Fiji software (to convert them into TIFF file format) and Imaris software for 3D image reconstruction. The number of intestinal stem cells (ISCs) and other cell counts in the gut were quantified using Imaris software (Bitplane).

#### Bacterial infections

*C. subtsugae ΔvioS* (*ΔvioS*) was grown in LB (Invitrogen) media inoculated with a single bacterial colony, taken from solid medium cultures grown from glycerol stocks kept at -80°C, and streaked fresh every week. All bacterial inocula were prepared from overnight liquid cultures, shaking at 30°C for 18 hours. The cultures were then diluted in phosphate buffer saline (PBS) to OD_600_=100. *Drosophila* adults were starved in empty fly vials for two hours at 29°C, after which they are added to food containing a filter soaked in a 1:1 mixture of 2.5% sucrose and the prepared bacterial suspension or LB media for controls. For each infection tube, 75µL of homogenous bacterial pellet mix (*OD_600_*=200) was mixed with an equal volume of 2.5% sucrose solution. In the non-infected control group, 75 µL of LB media was mixed with an equal volume of 2.5% sucrose solution. Flies were fed the inoculum for 24 hours and then were flipped onto new food or smurf food (normal food with FD&C Blue Dye 1 added for smurf assay). Infection vials with flies were maintained at 29°C, and death was recorded once a day for a number of days to monitor survival over time. For the Smurf assay, in addition to the number of deaths flies exhibiting systematic blue pigmentation were counted. Any flies that died without blue pigmentation were also noted.

#### Survival and smurf assays post-infection

On the day designated for the infection assay (Day 0), 30 two-day old female flies were sorted for each infection condition, which are placed into empty vials and incubated at 29°C for two hours and subsequently flipped into infection tubes. Dead flies recorded within this window of time are censored as infection-independent deaths.

For survival assay, following a 24-hour infection period, flies were transferred into vials containing filter paper soaked with 200uL 2.5% sucrose solution. The number of deaths was recorded every 24 hours, and flies were flipped into vials with fresh 2.5% sucrose every two days.

For smurf assay, following a 24-hour infection period, flies were transferred into vials containing smurf food. The number of dead flies was recorded and the flies were flipped into vials with fresh smurf food every 24 hours. Notably, the bodies of most deceased flies exhibited systematic blue pigmentation. Any flies that died without blue pigmentation were also noted.

#### Genetic ablation and regeneration assay in *Drosophila* male germline

GSCs and early-stage germ cells (up to 4-cell spermatogonia) were depleted using genetic manipulation of the pro-apoptotic gene, *grim*. The *UAS-grim* flies were crossed with *nanos-Gal4ΔVP16; tubulin-Gal80^ts^*and grown at 18°C to prevent *grim* expression, which is only turned on when shifting flies to 31°C to inactivate Gal80. The *nanos-Gal4ΔVP16; tubulin-Gal80^ts^*>*UAS-grim* flies were kept at 31°C for four days, which ablates early-stage germ cells (Abl in Fig. 4H, 4J). After 4-day ablation, flies were recovered at 18°C for another four days, when *grim* expression was inhibited again (Recv in Fig. 4H, 4J). After recovery, all GSCs including dedifferentiated GSC-like cells and *bona fide* GSCs were evaluated by immunostaining using anti-γ-Tubulin as the centrosome marker (*46*) and anti-Stat92E as the stemness marker (*54–57*).

#### Centrosome orientation assay

During aging or regeneration, both dedifferentiated GSC-like cells and *bona fide* GSCs were evaluated using centrosome orientation as a criterion, as shown previously (*18*) and illustrated (Fig. 4I, purple dot = centrosome). Centrosomes were immunostained using anti-γ-Tubulin(*46*). Centrosome misorientation is defined as neither of the two centrosomes being within the 90° hub–GSC interface (red in Fig. 4I). Centrosomes were scored to be oriented when one of two centrosomes is within the 90° hub–GSC interface (Fig. 4I).

#### *Drosophila* fertility assays

*polα50* P-element insertion flies were maintained over a balancer on the 3^rd^ chromosome. The *polα50^+/-^* flies used for fertility assays are from outcrossing the *polα50*/*Balancer* stock with *y,w* flies at 25°C to generate the F1 progeny, where the *polα50* P-element insertion allele is over a wild-type chromosome. As a control, *y,w* flies were crossed to another *wild-type* strain Oregon R flies. The F1 progeny for both *polα50^+/-^* and control were collected on the day they eclosed and aged simultaneously in separate vials.

For male fertility assays, the F1 *polα50^+/-^* or control males were aged at 25°C for the mentioned period of time with females. Vials were flipped every five days to prevent mixture with F2 progeny. When male flies (*polα50^+/-^* and *control^+/+^*) reached the desired age, one male was put into a new vial with three virgin *y,w* females. These flies were allowed to mate for five days, and then the parents were tossed away. Only those vials with all four parents (1 male and 3 female) alive after the 5-day mating period were retained for fertility assay. If any of the four parental flies died during the 5-day mating period, those vials were excluded from data collection and analyses. All F2 progeny were counted for 17-18 days after tossing the parent flies. Each mating from one male with three *y,w* females was recorded as one data point.

For female fertility assays, the F1 *polα50^+/-^* of control females were isolated as virgins and aged at 25°C in the absence of males to prevent mating before the assay. When female flies reached the desired age, one female was put into a new vial with two male *y,w* flies that had eclosed less than 24 hours previously. These flies were allowed to mate for five days, and then the parents were tossed away. Similar to the male fertility assay, only vials in which all parents survived the 5-day mating period were retained for the assay. All F2 progeny were counted for 15 days after tossing the parent flies. Each mating from one female with two *y,w* males was recorded as one data point.

#### *C. elegans* fertility assays

Manually selected L4 animals were grown individually on petri dishes seeded with OP50 *E. coli* food. They were then transferred on a new plate every 24 hours). The brood size of each worm was scored by counting the total number of larvae laid on the plates. For each brood size experiment, at least 30 worms were scored for each strain.

For hermaphrodite brood assays, the VC4505 strain is a CRISPR/Cas9 gene deletion of *pola-1*[*gk5576)III*] and maintained as a heterozygous strain. *pola-1(gk5576)/+* heterozygotes were used for experiments (*75*). As a control, the F1 progeny of both *pola-1(gk5576)* and the wild-type strain, N2, were scored in parallel on the same days.

Analysis of *C. elegans* germline GSC/progenitor zone

The strain GC1413 rrf-1(pk1417); naSi2 (Pmex-5::H2B::mCherry::nos-2 3′UTR); teIs113 (Ppie-1::GFP::H2B::zif-1 3′UTR) was used to label all germline nuclei with mCherry (red), while progenitor zone nuclei are doubly marked with GFP and mCherry (yellow) in both wild-type and *pola-1(gk5576)/+* heterozygote backgrounds. Quantification of each region was measured by counting the rows of cells from the distal end.

#### *Drosophila* lifespan assay

To assess the lifespan of *polα50^+/-^* and control flies, we adapted a protocol from (*73*). Groups of 20 newly eclosed flies (10 males and 10 females) were placed in vials with fly food and yeast. The day the flies eclosed was considered Day 0, and flies were kept in a temperature-controlled 25°C incubator for their entire lifespan. Every 2-3 days, flies were flipped onto fresh fly food with dry yeast, and deceased flies were removed and counted. Prior to Day 30, if all the males or all the females in a vial died (leaving all remaining flies as the same sex), then new flies of the missing sex were added, but not counted in the lifespan assay, to maintain the effects of mating in the remaining flies. Deceased flies were counted with respect to sex, but no significant difference was observed for control *versus* experimental males or control *versus* experimental females (data not shown).

#### *C. elegans* lifespan assays

All strains were maintained at 20°C. For each strain, gravid adults were bleached to isolate embryos. Embryos were then placed in liquid overnight to obtain a synchronized population of L1 worms that were then plated and allowed to grow to young adults. Strains N2 (n=68), CB1370 (n=33), and VC4505 (n=59), were transferred onto freshly seeded plates and scored by gently tapping with a platinum wire every 2-3 days.

#### Generation of human induced pluripotent stem cells with PolA1

To generate iPSC lines from human dermal fibroblast cells, the cells before five passages were used for the reprogramming experiments. Prior the CytoTune™-iPS 2.0 Sendai transfection, plate fibroblasts onto 6-well plates at ∼40% density with 2×10^5^– 3×10^5^ cells per well. During the early reprogramming stage, the 10-20 nM PolA1 inhibitor was added to the fibroblast growth medium: FBM^TM^ Basal Medium with FGM^TM^-2 SingleQuots^TM^ supplements (Lonza, CC-3131) until Day 7 and the fibroblast growth medium with PolA1 needs to be replaced daily. After that, the cells were passaged onto the vitronectin-coated dishes with 5×10^4^ cells per well and incubated 24 hours in a 37°C incubator with a humidified atmosphere of 5% CO2. Then change the medium to Essential 8™ Medium (Thermo Fisher, A1517001) to maintain the growth of programmed cells. The E8 medium needs to be replaced daily. Until Day 21, the colonies should have grown to an appropriate size for transfer. For the first three passages, manually cut the iPSCs colonies into small pieces and transfer onto a new vitronectin-coated dish. Allow the colonies to attach for 48 hours before replacing the spent medium with fresh E8 medium. When the colonies cover ∼80% of the surface area, passage the colonies using 0.5 mM EDTA prepared in Dulbecco’s Phosphate-Buffered Saline (DPBS) without calcium or magnesium. 10 μM Y-27632 was added to the E8 medium for one day for every passage.

#### Single-nucleus RNA-seq and data analysis

Colonies of induced human pluripotent stem cells (iPSCs) were picked and cultured in E8 medium until passage 9 at which time point single nuclei were isolated following 10× Genomics Protocol CG000365. One iPSC line with PolA1 inhibitor treatment and one without treatment (control) were used for experiments and data analysis.

Single-nucleus RNA-seq (snRNA-seq) libraries were prepared following 10x Genomics product instructions (Catalog# PN-1000283). The final libraries were sequenced by Novogene USA with paired-end 150 bp setting. The raw sequencing results were examined by fastqc v0.12.1 (https://www.bioinformatics.babraham.ac.uk/projects/fastqc/) before processing by cellranger v2.0.2 with GRCh38 as the reference genome. Publicly available single-cell RNA-seq datasets of human iPSCs were downloaded from ArrayExpress (accession number: E-MTAB-6524) (*101*) and Gene Expression Omnibus (accession number: GSE197380) (*102*) and used for comparison. RNA-seq count matrices of all samples were analyzed by Seurat v5.1.0 (*118*) after removing doublets with scDBlFinder v1.14.0 (*119*). The “CCAIntegration” method in Seurat was used to remove batch effects among samples before visualization in UMAP. The newly generated snRNA-seq data of iPSCs with or without PolA1 inhibitor treatment during induction were compared with published iPSCs scRNA-seq data, visualized in the same plot or separate plots.

## Supplemental Figures and Figure Legends

**Figure S1:**
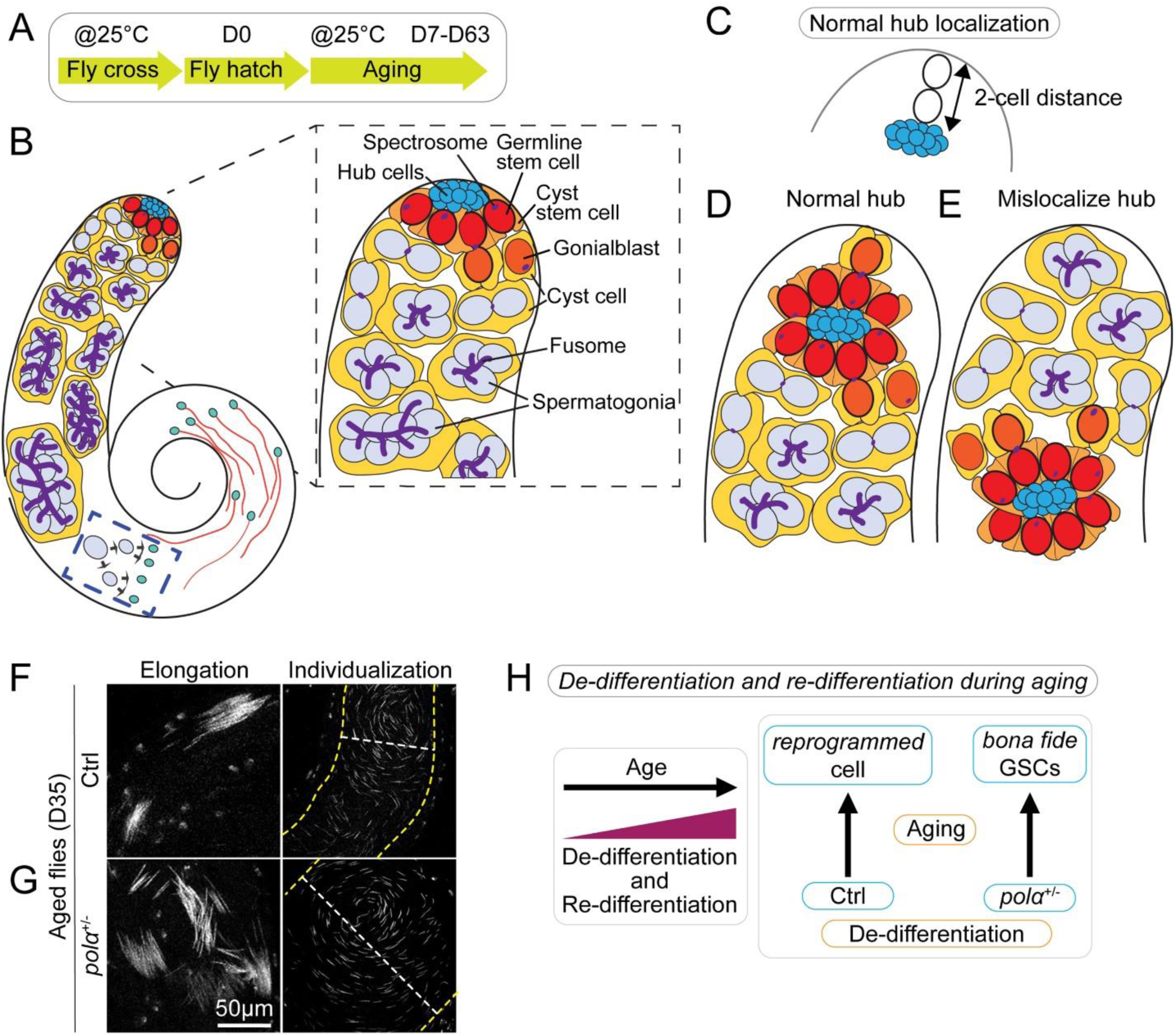
Design and results of the fertility assay in males with reduced Polα levels. (**A**) Regime of aging adult male flies at 25°C after eclosion (D0) up to 63 days (D63, see Materials and Methods). (**B**) A cartoon depicting a *Drosophila* testis and the apical tip showing different cell types and their characteristic cellular features. (**C-D**) A cartoon illustrating the apical tip of the testis with normal niche anatomy (**D**) and the criterion for normal hub localization (**C**). (**E**) A cartoon illustrating abnormal niche anatomy, with the hub structure mislocalized toward the middle of the testis. (**F-G**) DAPI staining of the spermatid elongation and individualization regions of the control (**F**) and *polα50^+/-^* (**G**) flies at D35 (35 days after eclosion). Scale bar: 50μm. The yellow dotted lines outline the testis in and white dotted line indicates the width of the basal part of testis. (**H**) A scheme showing the dedifferentiation and redifferentiation processes during aging.

**Figure S2:**
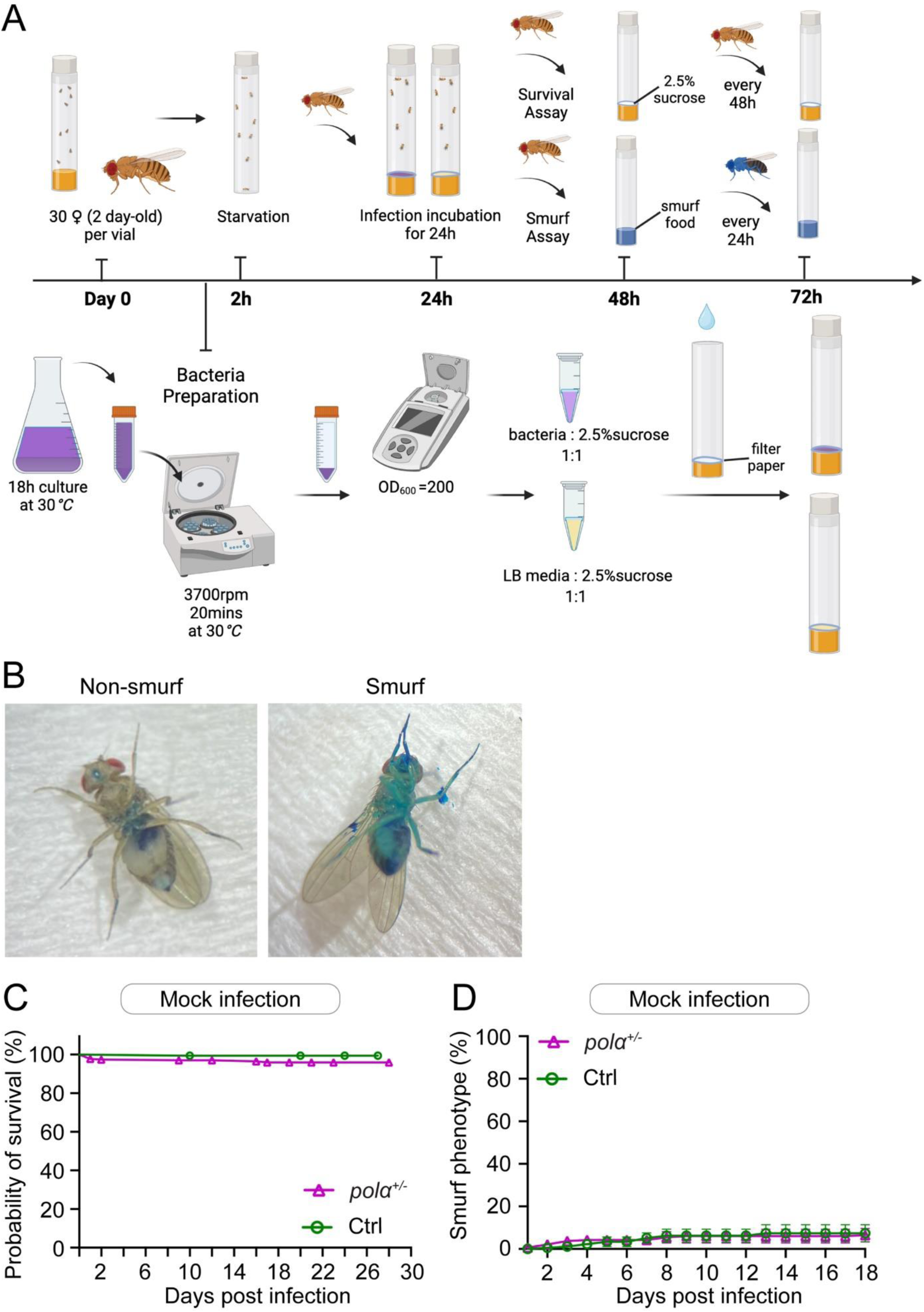
Experimental scheme and control experiments for survival and smurf assays post bacterial infection in *Drosophila* intestine. (**A**) Experimental scheme for bacterial infection inoculation, survival and smurf assays (see Methods and Materials for details). (**B**) Images showing a fly without smurf phenotype and a fly with smurf phenotype. (**C**) Probability of survival for non-infected controls: Flies were fed with a solution of LB media and sucrose as a mock infection for 24 hours, then replaced with fresh food every 48 hours: *pola50^+/-^*heterozygotes (magenta line, n= 271) *vs.* the control (green line, n = 178), *P*> 0.05 by Kaplan-Meier test: Log-rank (Mantel-Cox) test. (**D**) Smurf assay for non-infected controls: Using the same setting for mock infection as shown in (**C**), flies were flipped onto blue dye solid food daily to monitor the number of flies exhibiting smurf phenotypes: *pola50^+/-^*heterozygotes (magenta line, n = 209) *vs.* the control (green line, n =178). Two-way ANOVA (mixed-effects), for time factor: *P*=0.05; for genotype factor: *P*= 0.98.

**Figure S3:**
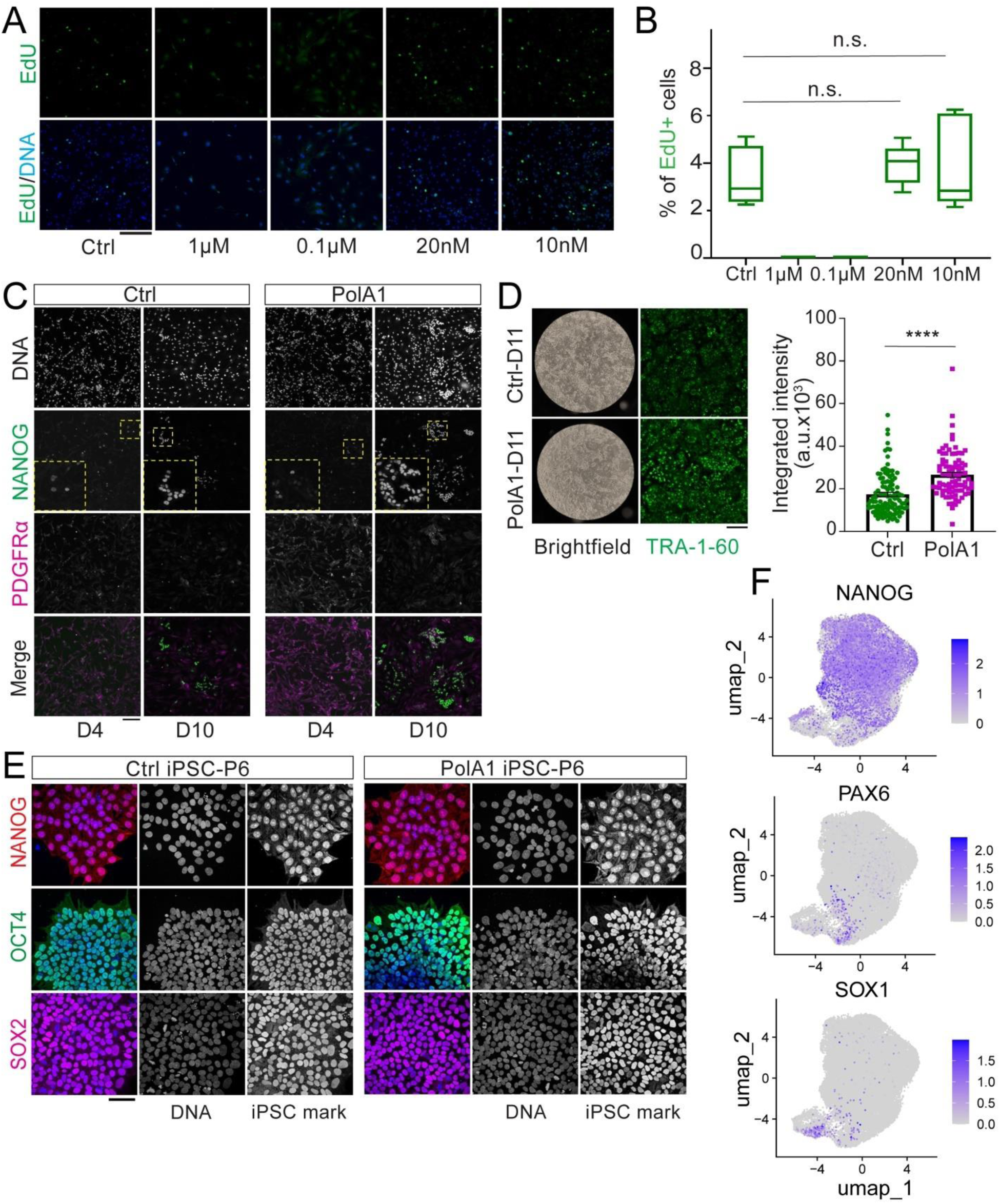
**Conditions and resultant cells from PolA1 inhibitor treated human dermal fibroblast cells in iPSC reprogramming process**. (**A**) EdU was pulsed for 20 minutes followed by quantification of EdU+ cells in human dermal fibroblast cells without and with different concentrations of PolA1 inhibitor. (**B**) Compared to the control (4.28±1.56%), no EdU+ cells were detected with PolA1 inhibitor treatment at high concentrations (10 µM, 1 µM, and 0.1 µM). In contrast, low concentrations of PolA1 treatment (0.02 µM: 4.95±1.09%, and 0.01 µM: 4.95±2.45%) show similar percentages of EdU+ cells: *P*=0.5368 (0.02 µM treatment *vs.* control) and *P*=0.8413 (0.01 µM treatment *vs.* control). All values = Average ±SEM, statistics was done using Mann Whitney test. (**C**) The NANOG and PDGFRα staining and at D4 and D10 during human iPSC reprogramming. (**D**) The live staining of TRA 1-60 in cells at D11 during reprogramming with and without PolA1 inhibitor. Ctrl: 17424±949.4; PolA1 treated: 26602±1137. \*\*\*\**P* <10^-4^, Average ± SEM by Mann Whitney test. (**E**) Immunofluorescence staining of iPSCs (Passage 6) shows strong expression of pluripotency markers in both control and PolA1 inhibitor treated human iPSCs. The pluripotency markers NANOG, OCT4 and SOX2 were examined using immunostaining. Scale: 50 µm. (**F**) Feature plots show expression of NANOG, a well-known pluripotency stem cell marker in all samples. Two differentiation markers, PAX6 and SOX1, are detected in a very small subset of cells, due to a technical issue with high confluence of the cells (*120*).

